# 4-Hydroxybenzaldehyde Attenuates Iso proterenol-induced Cardiac Fibrosis via TGF-β-Smad2/3 Signaling Pathway

**DOI:** 10.64898/2026.08.20.746132

**Authors:** Zhongcao Liu, Wei He, Fanglan Liu, Haifei Mao, Jian Chen

**Affiliations:** Shangrao Affiliated Hospital, Jiangxi Medical College, Nanchang University; Jiangxi Medical College, Shangrao, China

## Abstract

This study aims to investigate the cardioprotective effects of 4-Hydroxybenzaldehyde (4-HBA) against isoproterenol (ISO)-induced cardiac fibrosis and to elucidate the underlying mechanisms. In vivo, cardiac fibrosis was induced in C57BL/6 mice by subcutaneous injection of ISO, and the mice were treated with 4-HBA or a TGF-β inhibitor. Assessments using echocardiography, histopathology, and Western blotting demonstrated that 4-HBA significantly alleviated ISO-induced cardiac dysfunction, reduced collagen deposition, and attenuated apoptosis in mice. Mechanistically, 4-HBA inhibited TGF-β expression and Smad2/3 phosphorylation. In vitro, ISO was applied to cardiomyocytes (HL-1) and cardiac fibroblasts (MCFs), with or without 4-HBA or TGF-β inhibitor intervention. The results showed that 4-HBA suppressed HL-1 apoptosis and fibroblast proliferation, and significantly reduced the expression of extracellular matrix genes, TGF-β levels, and Smad2/3 phosphorylation in MCFs. These findings indicate that 4-HBA reduces myocardial injury while targeting the TGF-β/Smad2/3 pathway to attenuate cardiac fibrosis, highlighting its potential as a therapeutic agent for fibrotic cardiomyopathy.

## Introduction

Cardiovascular diseases (CVDs) are a leading cause of mortality and a major threat to human health worldwide [1, 2]. Cardiac fibrosis is a common pathological condition characterized by excessive accumulation of extracellular matrix (ECM), primarily collagen, in cardiac tissue [3]. This maladaptive remodeling increases the risk of arrhythmias and heart failure [4]. Recent advances have begun to elucidate the complex interactions among cardiac fibroblasts, cardiomyocytes, and the ECM during cardiac disease, identifying key pathways such as transforming growth factor-β (TGF-β) and SMAD2/3 that regulate fibroblast function and fibrotic progression [5, 6]. Isoproterenol (ISO) is a β-adrenergic agonist; overactivation of β-adrenergic receptors (β-ARs) readily triggers acute cardiac inflammation, myocardial injury, myocardial ischemia, and cardiac remodeling [8], and cardiac fibrosis is an adaptive response following β-AR activation [7, 9]. However, early use of beta-blockers has no significant association with in-hospital mortality in patients with stress cardiomyopathy [10]. Therefore, there is an urgent need to develop effective interventions that can halt ISO-induced cardiac injury and the progression of fibrosis. 4-Hydroxybenzaldehyde (4-HBA) is a naturally occurring phenolic aldehyde compound primarily found in the Chinese medicinal herb Gastrodia elata (Tianma) [11]. 4-HBA is most commonly used for the treatment of hypertension and exhibits a variety of biological activities, including antioxidant, anti-inflammatory, and vasorelaxant effects. 4-HBA is most commonly used in the treatment of hypertension and exhibits multiple biological activities, including antioxidant, anti-inflammatory, and vasorelaxant effects [12-15].

TGF-β plays a critical role in the pathological process of cardiac fibrosis [16]. The fibrotic cascade begins with TGF-β binding to its receptors on the surface of fibroblasts, initiating a Smad protein signaling cascade dominated by Smad2 and Smad3 [17]. Binding of TGF-β to its receptors induces receptor phosphorylation, which in turn activates Smad2 and Smad3. These phosphorylated proteins subsequently form heterodimeric complexes with Smad4 [18]. They translocate into the nucleus, where the heterodimeric complex regulates genes that control extracellular matrix (ECM) expression. Dysregulation of this pathway leads to excessive ECM accumulation, resulting in tissue stiffness and impaired organ function.

Herein, we validated in vivo that 4-HBA alleviates ISO-induced cardiac dysfunction and cardiac fibrosis in mice; In vivo and in vitro experiments demonstrated that 4-HBA not only protects cardiomyocytes but also has the potential to prevent fibroblast activation and excessive ECM deposition through the TGF-β/Smad2/3 pathway. These data support the TGF-β/Smad2/3 axis as a therapeutically targetable node through which 4-HBA exerts its anti-fibrotic and tissue-protective effects.

## Material and method

### Reagents and antibodies

Isoprenaline Hydrochloride (ISO), molecular formula C11H18ClNO3 (Cat# I8480), was purchased from Solarbio (Beijing, China). 4-Hydroxybenzoic acid(Cat#240141)was purchased from MERCK (Shanghai, China). Primary antibodies such as anti-ColⅠ (Cat# GB11022), anti-Col Ⅲ (Cat# GB151629), anti-α-Smooth Muscle Actin (α-SMA, Cat# GB111364), anti-Vimentin (Cat# GB11192), anti-Bax (Cat# GB114122), anti-Bcl-2 (Cat# GB154380) and anti-GAPDH (Cat# GB11002) were purchased from Servicebio (Wuhan, China). Anti-TGFβ (Cat# PA2154), anti-Phospho-Smad2/3 (Thr8, Cat# TA3367) and anti-Smad2/3 (Cat# PA1707) was purchased from Abmart (Shanghai, China) and anti-Mouse recombinant secondary antibody (Cat# RGAM001) and anti-Mouse recombinant secondary antibody (Cat# RGAM001) were purchased from Proteintech (Wuhan, China). TGF-β inhibitor SB-431542 (Cat# HY-10431R) was purchased from MCE (USA).

### Mice and animal models

Male specific-pathogen-free (SPF) C57BL mice, aged 6 to 8 weeks old, were acquired from Crisbio Biotechnology Co., Ltd. (Shanghai, China). The mice were housed in an environment maintained at a temperature ranging from 20-22 ℃, a relative humidity of 40%-60%, and a 12-hour light cycle.

Mice were randomly divided into four groups (n=6 per group) for a 14-day inject period. The control group received 100 μL isotonic saline (0.9% NaCl) via intraperitoneal injection (i.p.) every day. The ISO injected group was administered ISO (20 mg/kg body weight) dissolved in isotonic saline on the same schedule. For the 4-Hydroxybenzoic acid group (4-HBA), 4-HBA was first dissolved into isotonic saline at 20 mg/mL and then diluted with isotonic saline to achieve a dosage of 100 mg/kg body weight, 4-HBA was administered to the 4-HBA group via gavage over a period. The combination group received intraperitoneal injections of isoproterenol (ISO) at a dose of 20 mg/kg, while 4-hydroxybenzoic acid (4-HBA) was administered orally. All i.p. injections (saline, ISO) were administered at a volume 100 μL every day for 14 days.

At the end of experiments, the mice were anesthetized with 1% isoflurane, transthoracic echocardiography was evaluated using the Vevo 3100 high-resolution imaging system (VisualSonics, Canada) with a 30 MHz transducer. Following hair removal and application of ultrasound gel, two-dimensional B-mode and M-mode images were acquired from parasternal long-axis and short-axis views. Three consecutive cardiac cycles were recorded for each view. Left ventricular functional parameters including ejection fraction (EF), fractional shortening (FS) and stroke volume (SV) were measured offline using Vevo LAB software. After measurements, the mice were euthanized to collect the heart tissues. All experimental procedures were in compliance with the Guide for the Care and Use of Laboratory Animals.

### Cell culture and treatment

The mouse cardiomyocyte cell line HL-1 and Embryonic fibroblast cell line NIH/3T3 were obtained from the Chinese Academy of Sciences (Shanghai, China). Both cell lines were cultured in Dulbecco’s Modified Eagle Medium (DMEM, Servicebio, Wuhan, China) supplemented with 10% fetal bovine serum (FBS, Invitrogen, Carlsbad, CA, USA) in a humidified atmosphere with 5% CO_2_ at 37 °C. HL-1 cells were pretreated with ISO (10 μM), 4-HBA (10 μM), 4-HBA (10 μM) + ISO (10 μM), respectively for 6 hours, then replaced with fresh complete medium for additional 18 hours.

### Histopathological analysis

The heart tissues were initially fixed in 10% formaldehyde for 24 h, processed through dehydration, permeabilization, wax infiltration, and paraffin embedding, then sectioned into 4-μm slices. For histological analysis, sections were stained with hematoxylin and eosin (H&E, Beyotime Biotechnology) for general morphology assessment, Masson’s trichrome (Beyotime Biotechnology) for collagen fiber visualization, and Sirius red (Beyotime Biotechnology) for collagen quantification under polarized light, following the manufacturers’ protocols. Stained sections were imaged using a Nikon microscope. For quantitative analysis: (i) nuclear-cytoplasmic ratios were determined from H&E-stained sections by measuring 50 randomly selected cardiomyocytes per sample using ImageJ software (nuclear area/cytoplasmic area); (ii) fibrosis area percentage was calculated from Masson’s trichrome-stained and Sirius red-stained sections by thresholding blue-stained or red-stained collagen fibers in five random fields per sample.

The expression levels of BAX, Bcl-2 in the cardiac tissues were evaluated by immunohistochemical (IHC) analysis. IHC analysis was performed on 4-μm paraffin-embedded heart tissue sections using standard protocols. Briefly, the sections were treated with citrate (10 mM, pH = 6) for antigen retrieval, followed by counterstaining with Mayer’s hematoxylin for 15 min at room temperature. The sections were incubated with 5% bovine serum albumin (BSA) for 30 min at 37°C, then incubated with the following primary antibodies: rabbit anti-Bax (1:100) and rabbit anti-Bcl-2 (1:100) at 4°C overnight. After washing with PBS, sections were incubated with horseradish peroxidase (HRP)-conjugated goat anti-rabbit secondary antibody (1:100) for 30 min at 37°C. Finaly, the sections were then processed with 3,3′-diaminobenzidine (DAB) for 5 min at room temperature and counterstained with hematoxylin for 15 min at room temperature.

### TUNEL assay

Apoptotic cells in cardiac tissues were detected using the TUNEL Apoptosis Assay Kit (Beyotime Biotechnology) according to the manufacturer’s protocol. Briefly, after deparaffinization and antigen retrieval with Proteinase K (20 μg/mL, 15 min), the sections were incubated with TUNEL reaction mixture (60 min, 37°C), developed with DAB, and counterstained with hematoxylin. For the quantification of apoptosis, three random sections per sample were analyzed by counting TUNEL-positive nuclei in per section using ImageJ software. Results were expressed as percentage of TUNEL-positive cells (TUNEL⁺ nuclei/total nuclei ×100%).

### Western blot analysis

Total proteins were extracted from tissues or cells using RIPA mixture containing protease inhibitors and phosphatase inhibitors (Beyotime Biotechnology), followed by centrifugation at 12,000 g for 10 min at 4 ℃. Protein concentrations were determined by BCA assay (Beyotime Biotechnology), and approximately 20-30 μg of total protein were separated by 10% SDS-PAGE, and transferred onto polyvinylidene fluoride membranes. The membranes were blocked with 10% skim milk at room temperature for 2 h and washed with TBST for 10 min. They were then incubated at 4 ℃ overnight with primary antibodies against Col-Ⅰ (1:1000), Col-Ⅲ (1:1000), Vimentin (1:1000), ɑ-SMA (1:1000), Bax (1:1000), Bcl-2 (1:1000), ANP (1:1000), BNP (1:1000), TNF-α (1:1000), IL-1β (1:1000), TGF-β (1:1000), phospho-Smad2/3 (1:1000) and GAPDH (1:2000), followed by the secondary antibodies for 2 h. The Western blots were visualized using the ChemiDocTM XRS+ imaging system (Bio-Rad, California, USA). The expressions of the target proteins were normalized with GAPDH.

### Real-time quantitative RT–PCR (qRT-PCR)

qRT-PCR was conducted to assess the relative levels of mRNA expression. Total RNA was extracted from the samples using TRIzol reagent (Omega Bio-Tek, Norcross, Georgia, USA). After normalizing the RNA concentrations, the RNA was reverse transcribed into complementary DNA (cDNA) using a PrimeScript RT reagent kit (Takara, Beijing, China) with oligo(dT) primers. Subsequently, qRT-PCR analysis was conducted using the Power SYBR Green PCR Master Mix (vazyme Biotech Co.,Ltd, Nanjing, China). Relative mRNA expression levels of target genes were calculated using the 2^(-ΔΔCt) method with GAPDH as the endogenous control.

### Statistical analysis

Quantitative data were presented as means ± SEM and analyzed using GraphPad Prism 7.00 software. For comparisons between two groups, unpaired Student’s *t*-test was applied, and two-way ANOVA with Bonferroni post hoc test was used for multiple group comparisons. Statistical significance was considered as \**P* < 0.05, ** *P* < 0.01, *** *P* < 0.001, **** *P* < 0.0001.

## Result

### 4-HBA attenuates ISO-induced cardiomyocyte (HL-1) death and Embryonic fibroblast cell line (NIH/3T3) proliferation

To evaluate the potential cytotoxicity of 4-HBA on HL-1 cardiomyocytes and NIH/3T3, the CCK-8 assay was employed. Notably, when 4-HBA concentrations ranged from 10 to 100 µM, the optical density values of both HL-1 and NIH/3T3 were comparable to those of the normal control group. However, at a high concentration of 500 µM, significant cell death was observed in both cell types, indicating a cytotoxic threshold. These results demonstrate that 4-HBA at concentrations below 100 µM is non-toxic to HL-1 and NIH/3T3 (Fig. 1A–B). ISO-induced cardiac injury and fibrotic progression lead to cardiomyocyte death and abnormal proliferation of cardiac fibroblasts. When cells were co-treated with ISO and 4-HBA at 10 or 50 µM, the detrimental effects of ISO were markedly attenuated, suggesting that 4-HBA possesses the potential to protect cardiomyocytes and prevent excessive fibroblast proliferation (Fig. 1C–D).

**Fig 1.**
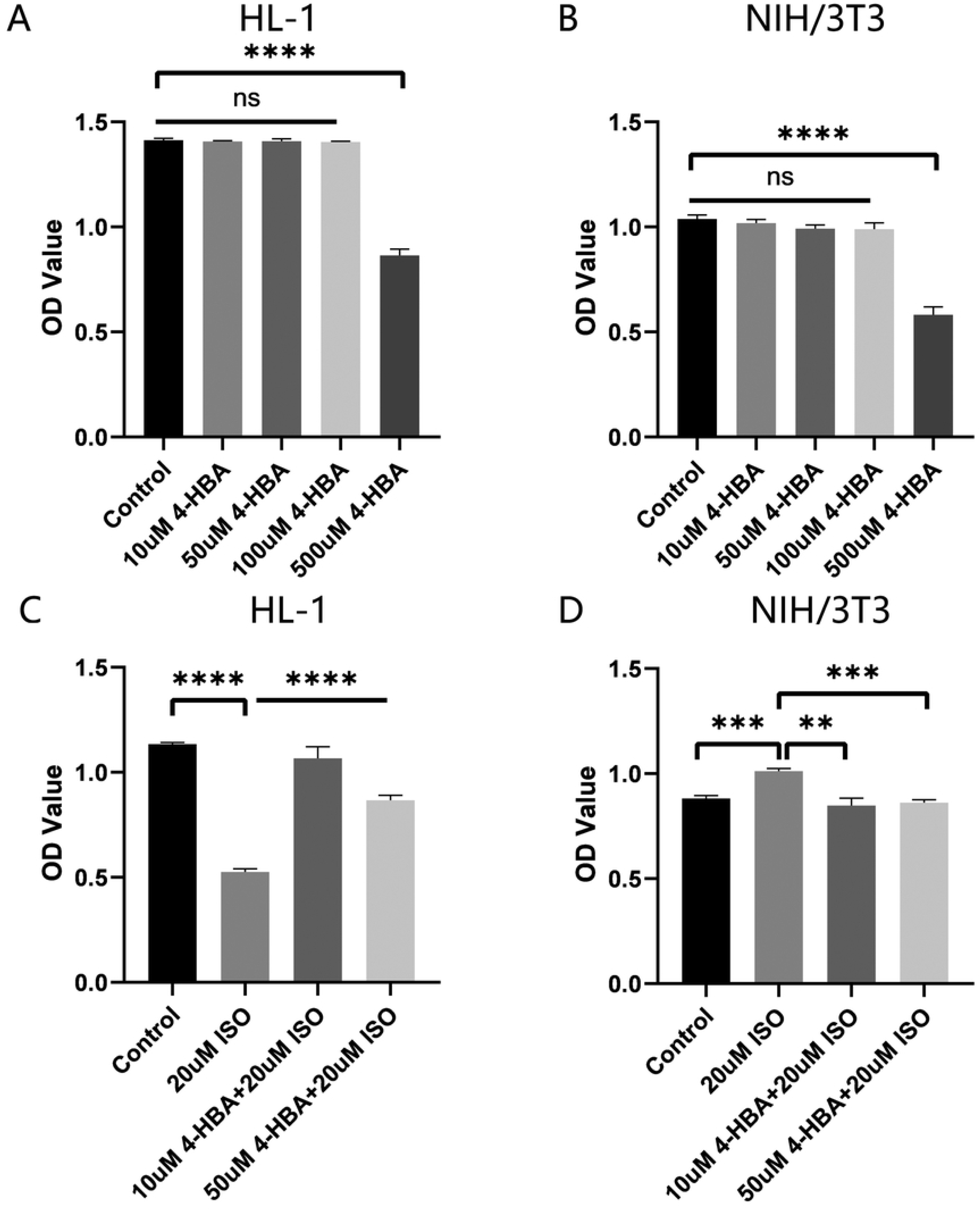
4-Hydroxybenzaldehyde (4-HBA) reduces the damaging effects of isoproterenol (ISO) on HL-1 and NIH/3T3 cells. **(A)** Cytotoxicity in HL-1 cells after 24 h treatment with various concentrations of 4-HBA (10, 50, 100, and 500 μg/mL). (**B**) Cytotoxicity in NIH/3T3 cells after 24 h treatment with various concentrations of 4-HBA (10, 50, 100, and 500 μg/mL). **(C)** Cell viability results of HL-1 cells in the NC, ISO, and ISO + 4-HBA groups. **(D)** Cell viability results of NIH/3T3 cells in the NC, ISO, and ISO + 4-HBA groups. NS P > 0.05, *P < 0.05, **P < 0.01, ***P < 0.001, ****P < 0.0001.

### 4-HBA inhibits ISO-promoted apoptosis in cardiomyocytes (HL-1)

In vitro experiments showed that co-administration of 10μM or 50μM 4-HBA and ISO significantly reduced the apoptotic rate of HL-1 cells. These results were further supported by flow cytometry and immunofluorescence staining (Fig. 2 A-D).

**Fig 2.**
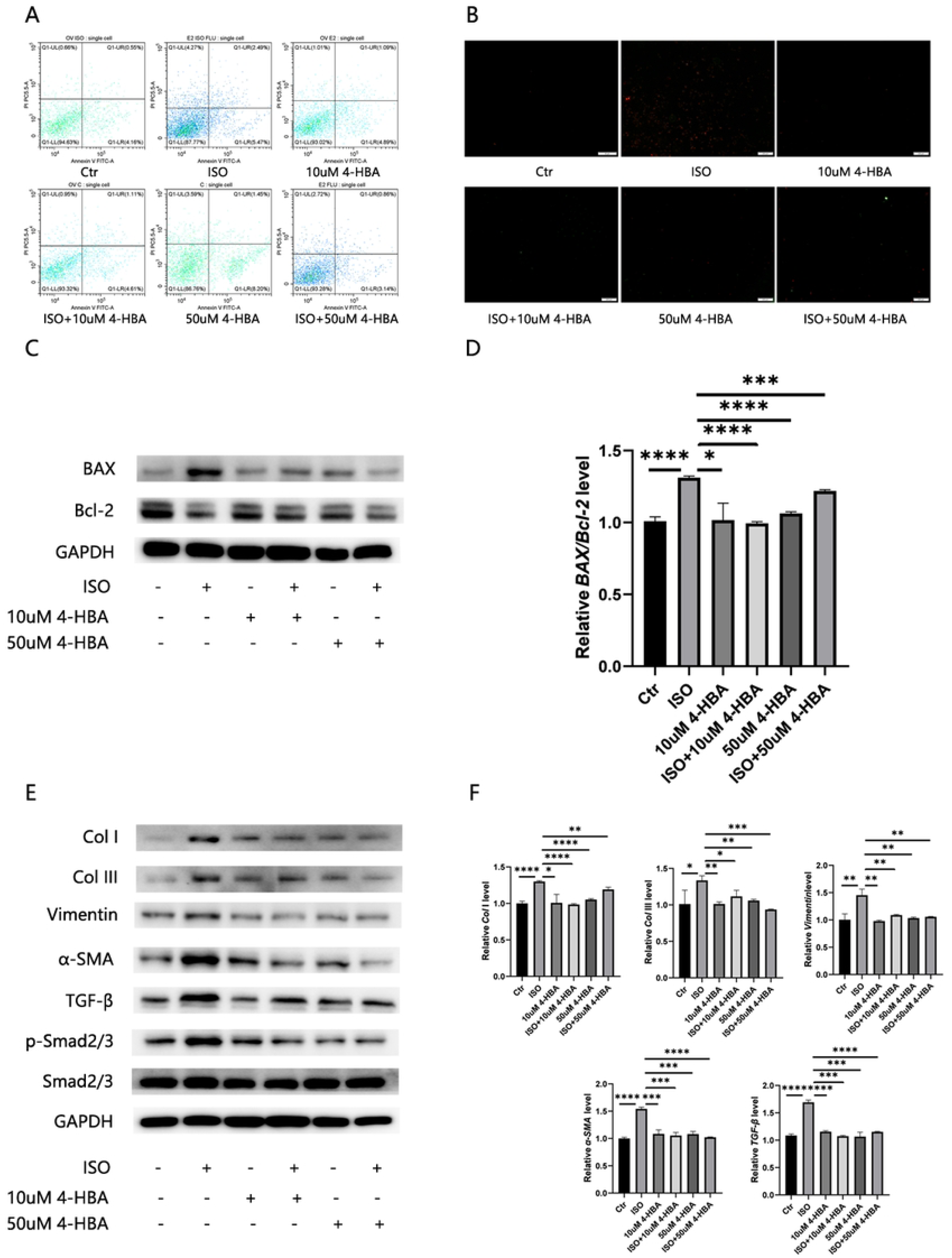
4-HBA suppresses ISO-induced apoptosis in HL-1 cells and pro-fibrotic effects in NIH/3T3 cells. **(A)** Apoptotic cells were identified using flow cytometry with Annexin V-FITC. **(B)** Apoptotic cells were detected using Annexin V-PI staining. (**C**) Western blot analysis of apoptosis-related proteins Bax and Bcl-2 in HL-1. **(D)** The mRNA expression levels of Bax/Bcl-2 ratio in HL-1. **(E)** Western blot analysis of fibrosis-related protein Col I, Col III, Vimentin, α-SMA, TGF-β and SMAD in NIH/3T3. **(F)** The mRNA expression levels of Col I, Col III, Vimentin, α-SMA and TGF-β in NIH/3T3. *P < 0.05, **P < 0.01, ***P < 0.001, ****P < 0.0001.

### 4-HBA suppresses ISO-induced pro-fibrotic effects in cardiac fibroblasts(MCFs)

In vitro experiments demonstrated that co-treatment with 10μM、50μM 4-HBA and ISO markedly decreased fibrotic markers in MCFs and modulated the pro-fibrotic TGF-β1/Smad signalling pathway. (Fig. 2 E-F).

### 4-HBA attenuates ISO-induced cardiac dysfunction, collagen deposition and apoptosis in mice

To elucidate the protective mechanism of 4-HBA against ISO-induced cardiac injury, mice were administered 4-HBA by oral gavage at a dose of 100 mg/kg/day for 14 days. After completion of the experimental protocol, echocardiographic assessment revealed that 4-HBA treatment restored ejection fraction, fractional shortening and stroke volume to near-normal levels (Fig. 3 A).

**Fig 3.**
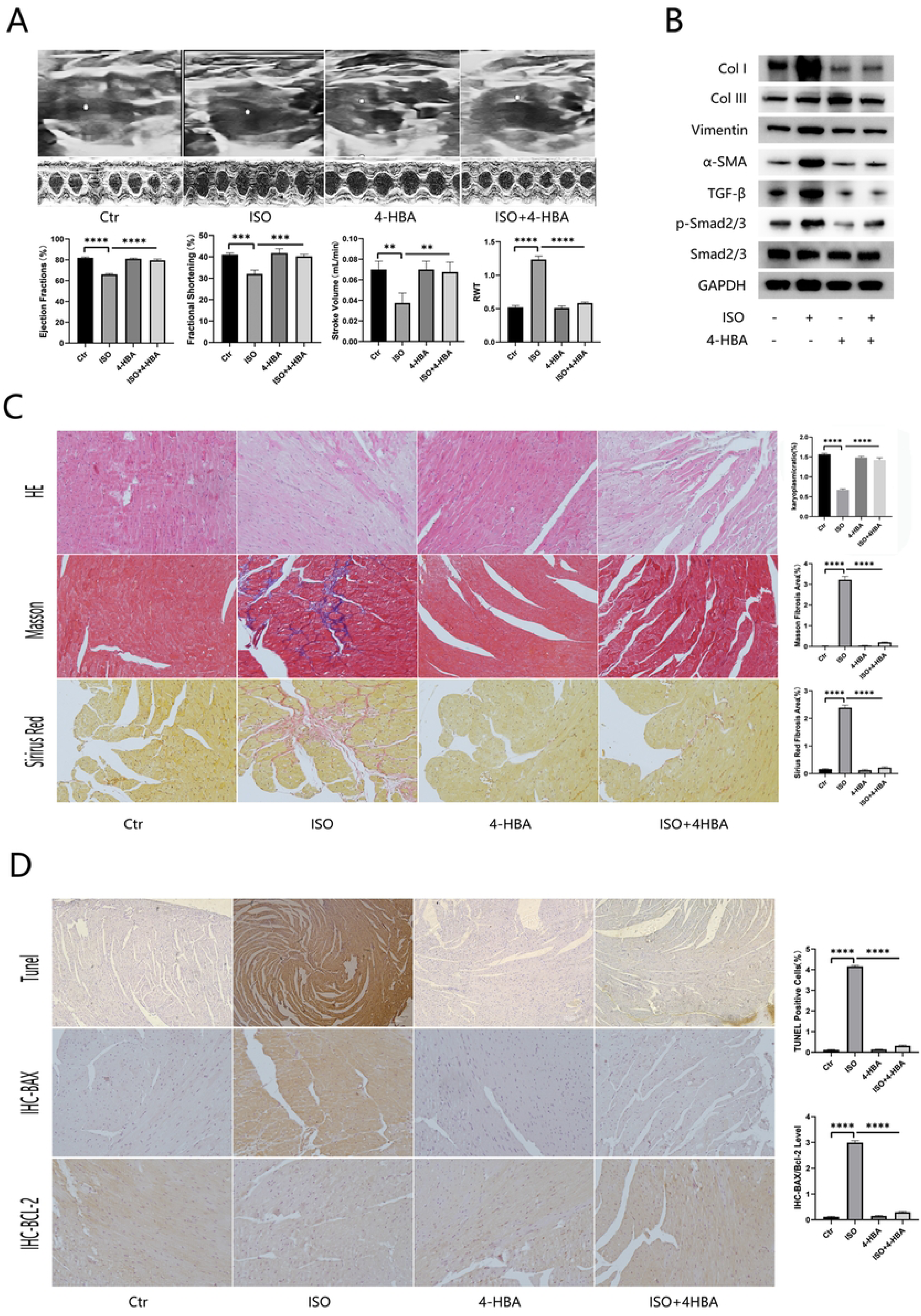
4-HBA ameliorates ISO-induced cardiac dysfunction in mice. **(A)** Representative images of echocardiography of mouse hearts and the average data of cardiac function of left ventricular EF, FS, SV and RWT. **(B)** Western blot analysis of fibrosis-related protein Col I, Col III, Vimentin, α-SMA, TGF-β and SMAD in cardiac tissue. **(C)** Representative images of HE, Masson and Sirius red staining of cardiac tissue and the quantification of nucleus-to-cytoplasm ratio and fibrosis area by Masson and Sirius red staining. **(D)** Representative images of TUNEL, Bax and Bcl-2 staining images of cardiac tissues. **(E)** Representative images of HE, Masson and Sirius red staining of cardiac tissue and the quantification of nucleus-to-cytoplasm ratio and fibrosis area by Masson and Sirius red staining. **(F)** Representative images of TUNEL, Bax and Bcl-2 staining images of cardiac tissues. *P < 0.05, **P < 0.01, ***P < 0.001, ****P < 0.0001.

**Fig 4.**
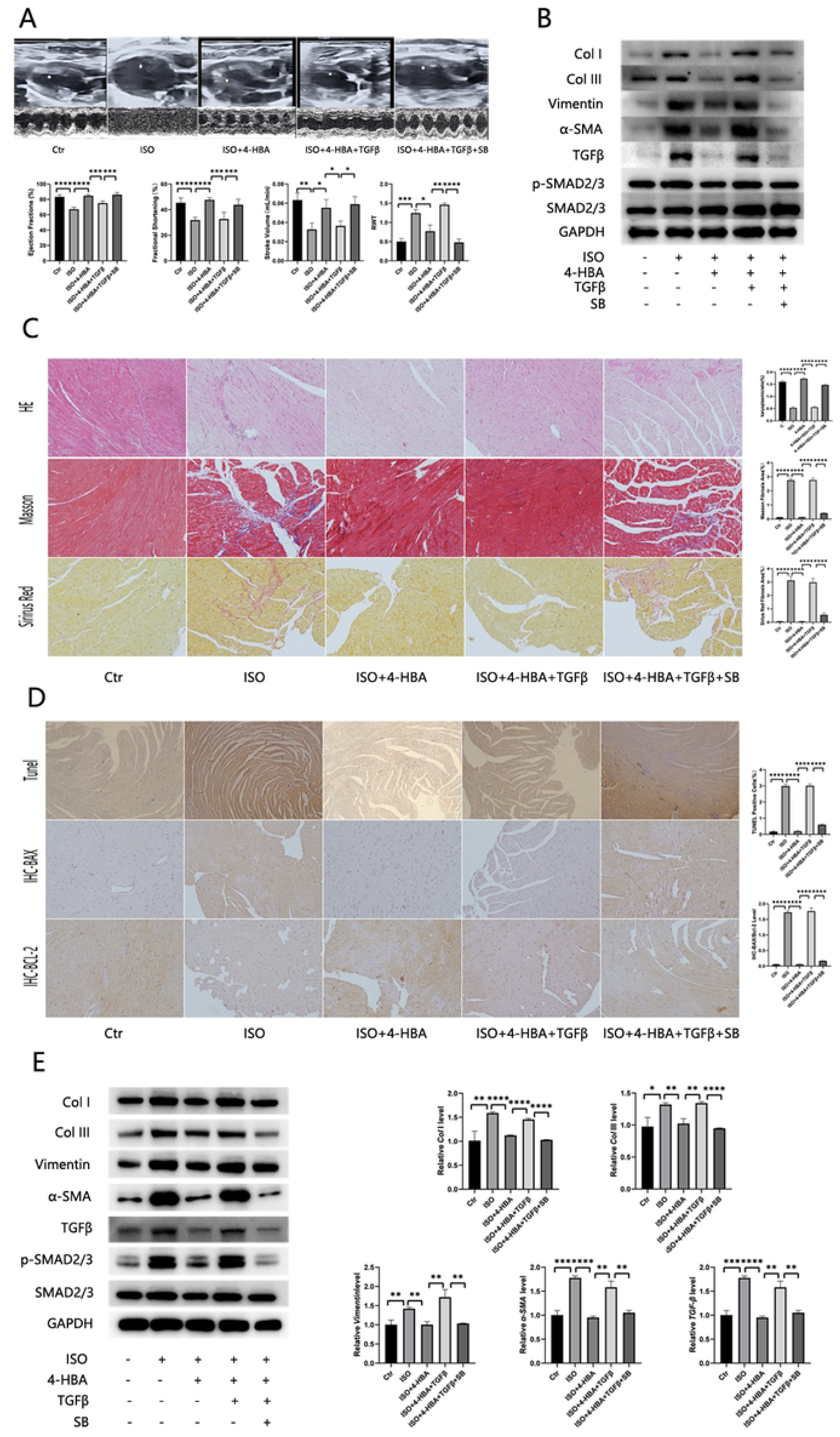
4-HBA attenuates ISO-induced cardiac fibrosis through the TGF-β1/Smad signaling pathway. **(A)** Representative images of echocardiography of mouse hearts and the average data of cardiac function of left ventricular EF, FS, SV and RWT. **(B)** Western blot analysis of fibrosis-related protein Col I, Col III, Vimentin, α-SMA, TGF-β and SMAD in cardiac tissue. **(C)** Representative images of HE, Masson and Sirius red staining of cardiac tissue and the quantification of nucleus-to-cytoplasm ratio and fibrosis area by Masson and Sirius red staining. **(D)** Representative images of TUNEL, Bax and Bcl-2 staining images of cardiac tissues. **(E)** Western blot analysis of fibrosis-related protein Col I, Col III, Vimentin, α-SMA, TGF-β and SMAD in NIH/3T3. **(F)** The mRNA expression levels of Col I, Col III, Vimentin, α-SMA and TGF-β in NIH/3T3. *P < 0.05, **P < 0.01, ***P < 0.001, ****P < 0.0001.

Histopathological and Western blot analyses further demonstrated that 4-HBA administration significantly reduced the area of myocardial fibrosis and downregulated the expression levels of fibrotic markers, including COL I/III, α-SMA and Vimentin, and modulated the pro-fibrotic TGF-β1/Smad pathway (Fig. 3 B-C). TUNEL staining showed that the apoptotic rate in mouse myocardial tissue returned to near-normal levels after 4-HBA administration (Fig. 3 D). Immunohistochemical analysis revealed that the expression levels of BAX and Bcl-2 were normalised after 4-HBA treatment (Fig. 3 D).

### 4-HBA is associated with TGF-β1/Smad signaling pathway modulation

The in vivo and in vitro results indicated that 4-HBA can modulate the TGF-β1/Smad pathway; however, whether it primarily reduces ISO-induced cardiac dysfunction and myocardial fibrosis through this pathway remained uncertain. Therefore, we explored this by administering a TGF-β1 inhibitor prior to 4-HBA treatment.

In vivo experiments demonstrated that TGF-β1 inhibitor treatment blocked the protective effects of 4-HBA against ISO-induced cardiac injury and inhibited its regulatory effects on TGF-β1 expression and Smad2/3 phosphorylation in cardiac tissue.

In vitro experiments showed that TGF-β1 inhibitor treatment prevented the effects of 4-HBA in reducing fibrotic markers, TGF-β1 expression and Smad2/3 phosphorylation levels in MCFs.

## Discussion

This study is the first to systematically elucidate the protective effects of 4-hydroxybenzaldehyde (4-HBA) against isoproterenol (ISO)-induced cardiac fibrosis and its underlying molecular mechanisms. Through both in vitro and in vivo experiments, we demonstrated that 4-HBA significantly attenuates ISO-induced cardiac dysfunction, cardiomyocyte apoptosis, and aberrant proliferation of cardiac fibroblasts, and inhibits excessive extracellular matrix deposition. The mechanism is closely associated with targeted modulation of the TGF-β/Smad2/3 signalling pathway.

The core pathological mechanism of cardiac fibrosis lies in the overactivation of the TGF-β signalling pathway [19]. Upon binding to its receptor, TGF-β phosphorylates Smad2/3, which then forms a complex with Smad4 and translocates to the nucleus to initiate the transcription of pro-fibrotic genes such as collagen (Col I/III) and α-SMA [20-21].In the present study, ISO treatment significantly elevated TGF-β expression and enhanced Smad2/3 phosphorylation both in vivo and in vitro, whereas 4-HBA intervention effectively reversed these changes. More importantly, we performed mechanistic validation using a TGF-β inhibitor: when TGF-β signalling was blocked, the protective effects of 4-HBA were abolished, directly indicating that the TGF-β/Smad2/3 pathway is the key target through which 4-HBA exerts its anti-fibrotic and cardioprotective actions.

At the cellular level, 4-HBA displayed dual regulatory properties. In cardiomyocytes (HL-1), 4-HBA significantly reduced the ISO-induced apoptotic rate, as confirmed by flow cytometry and immunofluorescence staining. In cardiac fibroblasts (MCFs), 4-HBA inhibited aberrant cell proliferation and the pro-fibrotic phenotypic transformation (e.g., downregulation of α-SMA and collagen expression). Notably, 4-HBA showed no cytotoxicity at concentrations of 10–100 µM, whereas toxicity appeared at 500 µM, indicating a relatively broad safety window and providing a dose reference for subsequent clinical translation. In the whole-animal model, oral gavage of 4-HBA (100 mg/kg/day for 14 days) effectively improved cardiac function (restoring ejection fraction, fractional shortening, and stroke volume) in ISO-treated mice, reduced myocardial collagen deposition and apoptosis, and restored the balance of Bcl-2/Bax. These results were highly consistent with the in vitro findings, further confirming the systemic protective effects of 4- HBA.

HBA is a natural phenolic compound mainly found in the traditional Chinese medicine Gastrodia elata [1].Previous studies have reported its antioxidant, anti-inflammatory, and vasorelaxant activities [11-14]. The novelty of this study lies in extending the application of 4-HBA to the field of cardiac fibrosis and revealing a new mechanism of its anti-fibrotic action via the TGF-β/Smad2/3 pathway. This provides a new candidate molecule for the development of natural- product-based anti-fibrotic drugs.

However, this study has certain limitations. First, although we confirmed that TGF-β/Smad2/3 is the major pathway, whether 4-HBA simultaneously affects other signalling pathways (e.g., non-Smad-dependent pathways, oxidative stress pathways, etc.) remains to be explored. Second, the oral bioavailability and metabolite activity of 4-HBA need further clarification. In addition, this study only observed short-term effects over 14 days; the long-term efficacy and safety of 4-HBA in chronic cardiac fibrosis still require validation in longer-term models.

## Conclusions

This study is the first to systematically demonstrate that 4-hydroxybenzaldehyde (4-HBA) significantly attenuates isoproterenol (ISO)-induced cardiac fibrosis and cardiac dysfunction. Both in vivo and in vitro experiments confirmed that 4-HBA alleviates ISO-induced cardiomyocyte apoptosis, suppresses abnormal proliferation of cardiac fibroblasts, and reduces excessive extracellular matrix deposition. Mechanistically, 4-HBA exerts its cardioprotective and anti-fibrotic effects primarily through targeted inhibition of the TGF-β/Smad2/3 signaling pathway. These findings indicate that 4-HBA holds promise as a potential therapeutic candidate for the treatment of fibrotic cardiomyopath. Future studies are warranted to further optimize the dosing strategy of 4-HBA, explore its long-term efficacy and safety, and investigate potential combination therapies targeting other anti-fibrotic pathways.

## Acknowledgment

The authors gratefully acknowledge DeepSeek V4.0 (www.deepseek.com) for providing English grammar correction during the preparation of this manuscript.

